# The Italian Autism Network (ITAN): A Resource for Molecular Genetics and Biomarker Investigations

**DOI:** 10.1101/306845

**Authors:** Pierandrea Muglia, Michele Filosi, Lucio Da Ros, Tony Kam-Thong, Franco Nardocci, Elisabetta Trabetti, Emiliangelo Ratti, Paolo Rizzini, Alessandro Zuddas, Bernardo Dalla Bernardina, Enrico Domenici, the Italian Autism Network

## Abstract

**Background:** A substantial genetic component accounts for Autism Spectrum Disorders (ASD) aetiology, with some rare and common genetic risk factors recently identified. Large collections of DNAs from thoroughly characterized ASD families are an essential step to confirm genetic risk factors, identify new variants and investigate genotype-phenotype correlations. The Italian Autism Network aimed at constituting a clinical database and a biorepository of samples derived from ASD subjects and first-degree relatives extensively and consistently characterized by child psychiatry centers in Italy.

**Methods:** The study was approved by the ethical committee of the University of Verona, the coordinating site, and by the local ethical committees of each recruiting site. Certified staff was specifically trained at each site for the overall study conduct, for clinical protocol administration and handling of biological material. A centralized database was developed to collect clinical assessment and medical records from each recruiting site. Children were eligible for recruitment based on the following inclusion criteria: age 4–18 years, at least one parent or legal guardian giving voluntary written consent, meeting DSM-IV criteria for Autistic Disorder or Asperger’s Disorder or Pervasive Developmental Disorder NOS. Affected individuals were assessed by full psychiatric, neurological and physical examination, evaluation with ADI-R and ADOS scales, cognitive assessment with Wechsler Intelligence Scale for Children or Preschool and Primary, Leiter International Performance Scale or Griffiths Mental Developmental Scale. Additional evaluations included language assessment, the Krug Asperger’s Disorder Index, and instrumental examination such as EEG and structural MRI. DNA, RNA and plasma were collected from eligible individuals and relatives. A central laboratory was established to host the biorepository, perform DNA and RNA extraction and lymphocytes immortalisation.

**Discussion:** The study has led to an extensive collection of biological samples associated with standardised clinical assessments from a network of expert clinicians and psychologists. Eighteen sites have received ADI/ADOS training, thirteen of which have been actively recruiting. The clinical database currently includes information on 812 individuals from 249 families, and the biorepository has samples for 98% of the subjects. This effort has generated a highly valuable resource for conducting clinical and genetic research of ASD, amenable to further expansion.

## Background

Autism Spectrum Disorders (ASD), also previously referred to as Pervasive Developmental Disorders (PDD), are a group of neurodevelopmental disorder with onset in early childhood characterized by a triad of core symptoms: impaired social interaction, compromised language and communication and repetitive and narrow pattern of behaviors and interests. Many neuropsychiatric conditions and symptoms are often present in addition to core symptoms in ASD; the most common being mental retardation, hyperactivity, epilepsy, aggressive behaviours. Autism or Autistic Disorder (AD), is the most common and well-known form of ASD that includes Asperger Syndrome (AS), Pervasive Developmental Disorder Not Otherwise Specified (PDD – NOS), Childhood Disintegrative Disorder (CDD) and Rett Syndrome (RS). All above subtypes are falling under the single diagnostic category of autism spectrum disorders, according to the new revision of the Diagnostic and Statistical Manual of Mental Disorders (DSM-V) [1]. AS is a less severe variant of AD, in which there is no cognitive deficit or language impairment. PDD - NOS defines a subtype of AD that characterizes subjects who are on the autism spectrum but do not fully meet the criteria for other AD or other ASD, and it was considered a separate diagnostic category in the DSM-IV. CDD and RS are more rare forms of ASD and show more distinct features from the rest of ASD [2]. CDD is a rare and not very well-known variant of ASD characterized by normal development followed by marked regression in many skills that begin after the age of two [3,4]. Rett Syndrome (RS) is caused in 80% of cases by sporadic mutations of *MECP2* located on Xq28. As opposed to AD, RS is more common in females. It has a lifetime prevalence of about 1 in 20.000 females and is characterized by apparently normal development for the first 6-18 months of life, followed by a period of regression in language and motor skills [5]. Both CDD and RS represent nosologically distinct disorders from AD, AS and PDD - NOS. In this manuscript, we use the term ASD to refer to AD, AS and PDD-NOS since the three diagnoses can be considered as a continuum rather than clear distinct disorders, consistently with the newly revised DSM-5. Within each of the three diagnostic categories, clinical heterogeneity is substantial. The fact that AD children share the same diagnostic categories does not imply they share set or severity of core and ASD-associated symptoms. Also, heterogeneity on ASD aetiology is more than likely. Several etiological hypotheses exist, such as altered synaptic dysfunction [6] leading to imbalanced excitatory and inhibitory neurotransmission [7]. However, a unifying aetiological theory is still lacking, and it is more than likely that the clinical heterogeneity is also paralleled by etiological heterogeneity.

Prevalence of ASD is reported to be much higher in most recent studies as compared to studies conducted a decade ago. Recent epidemiological studies indicate a worldwide prevalence for ASD ranging from 0.6 to 0.7 % with an approximate males to females ratio of 4-5 to 1 [8, 9], as compared to the prevalence of less than 0.2 % in earlier investigations [10]. The interpretation of increased prevalence detected in recent studies is not straightforward and has raised controversial debates on the potential causes. The original paper published in Lancet [11] that initiated the controversy on vaccines as a putative cause of autism has been recently retracted by the Journal since fraud has been detected. The vast majority of the scientific community agrees on the unlikelihood of vaccination as a cause. One confounding factor, when reporting prevalence estimates, could be the change over time of diagnostic criteria for ASD, making the comparisons of epidemiological studies more difficult. It is commonly believed that the increase in the overall reported prevalence is mostly due to the development of more comprehensive ascertainment and to the expansion of diagnostic categories, which include milder forms of the disorder [12]. Considerable evidence from twin studies indicates a substantial genetic component in the aetiology of ASD. ASD heritability has been estimated to be between 80 and 90% according to recent family and twin studies [13]. Monozygotic twin concordance has been found to as high as >88%, whilst in dizygotic pairs and siblings the concordance rates are >30% and >15%, respectively, higher than what previously reported, suggesting a significant environmental contribution to ASD [14]. Substantial heterogeneity also exists for the genetic component of ASD, where the interplay between commonly inherited variants (single nucleotide polymorphism or SNPs) and highly penetrant rare variations results with a complex genetic architecture [15]. A small proportion of ASD cases has been associated to *de novo*copy number variations (CNV), including deletions and duplications, as well as to *de novo* pathogenic single nucleotide variations (SNVs), accounting for a total of 3% of cases according to recent estimates [16]. Furthermore, ASD related syndromes for which the genetic cause is known account for additional 3.4% of ASD. These data support a polygenic model, where most ASD liability resides with many common variations along the genome, although the total genetic contribution to ASD (including *de novo* variations, syndromic mutations, non-additive and additive SNP effects) does not exceed 60%, with the remaining proportion being unaccounted for [16] Still, the increased number of rare variants exhibiting large individual effect sizes leaves open the single major gene model, consistent with earlier estimates suggesting about 10 to 20% of ASD cases to be due to known genetic abnormalities [17].

The most common ASD associated syndromes include: i) Fragile X syndrome (caused by expansion of polymorphic CGG repeat upstream of the FMR1 gene) in which over 40% show Autism core symptoms; ii) Angelman syndrome, determined by a deletion of 15q11-13 where ASD is present in about 50% of cases; iii) Tuberous Sclerosis, due to mutations in two functionally-related genes, TSC1 and TSC2, in which autism is present in more than 60% cases; iv) 22q deletion (or Phelan-McDermid) syndrome where about 80% of cases present with autistic symptoms [16].

Recent sequencing approaches (including exome-or whole-genome sequencing) have contributed to the identification (or validation) of highly penetrant rare genetic variants, including CHD8, GRIN2B, SCN2A, and SYNGAP1 [18] (see also [19] for a recent review). *De novo* mutations associated to ASD are often located in genes functionally correlated. Among them, neuroligin 3 and 4 (NLGN3 and NLGN4), Neurexin3, SHANK3 and CNTNAP2, which are all involved in synapse formation and specialization, supporting the notion that altered synaptic formation can lead to abnormal neural pathways in ASD) [6]. However, it is important to underline that no single neurobiological hypothesis yet can explain the aetiology of ASD, which is still elusive and likely to be determined by multiple genetic and non-genetic factors.

As far as the contribution from commonly inherited variants, the advent of affordable high-throughput genotyping technologies allowing for genome-wide association studies (GWAS), combined with the increased availability of ASD collections, is beginning to pay-off. Although limited in statistical power to robustly identify genome-wide significant loci, early studies have shown the potential of GWAS in the identification putative novel risk variants [20]. Two of the first GWAS in ASD reported the identification of common genetic variants residing between two genes mapping on the chromosome 5 and coding for cell molecule adhesion proteins (cadherin 9 and cadherin 10), providing support to a previously hypothesized altered neuronal cell adhesion in ASD [21,21]. A third study, based on a broad collaborative effort, lead to the identification of an additional common variant on chromosome 5, adjacent to several genes including *SEMA5A*, a member of semaphorin protein involved in axonal guidance [23]. Later, the Autism Genome Project (AGP, The AGP Consortium, representing more than 50 centers in North America and Europe) identified genome-wide association signals for SNPs located in the gene MACROD2 [24] and CNTNAP2, a gene previously implicated in ASD [25]. Most of the above findings were not replicated in a subsequent meta-analysis in European samples [26, 27], suggesting the need for larger ASD cohorts to clarify the role of common variation in the disorder. Large-scale coordinated international collaborations such as the Psychiatry Genomic Consortium (PGC) have been established, with the aim of conducting statistically rigorous and comprehensive GWAS meta-analyses for major psychiatric disorders [28, 29]. The first meta-analysis of the ASD Working Group of the Psychiatric Genomics Consortium, conducted on over 16.000 ASD individuals has been recently published [30]. The study did not show individual variants exceeding the accepted GWS threshold in the discovery dataset. However, it was able to identify a novel and genome-wide significant association on chromosome 10q24.32 (mapping in CUEDC2) in the combined metaanalysis and confirmed among top-ranking associations some of the findings from previous GWAS.

Notwithstanding the impact of modern genome-wide approaches in the dissection of the complex genetic component of ASD, the vast majority of it is still to be unveiled. The analysis of larger and, in particular, clinically well-characterized ASD samples is critical for the identification of new genetic variants and their validation, and for establishing of genotype-phenotype correlations. With these premises, the ‘Fondazione Smith Kline’ has supported the creation of an area for autism research that has launched the Italian Autism Network (ITAN) initiative. This initiative aims to create a repository of biological material (DNA, RNA and plasma) and a database of clinical information on autism individuals and their first-degree relatives enabling research on the aetiology and phenomenology of ASD. The recruitment of the first families began back in 2008. In this article, we describe the protocol used by all recruiting sites for clinical evaluation of the ASD subjects and relatives. We also provide the main clinical features of the 249 recruited families.

## Methods

### Study Design

The ‘Fondazione Smith Kline’ Italy, with its “Autism Research Area”, subsequently spun out as a separate Foundation (“ITAN - Italian Autism Network”), has sponsored a multi-centric project to recruit and assess autism individuals and their first-degree relatives to create a repository of genomic DNA, plasma, peripheral RNA and lymphoblastoid cell lines (LCLs). Extensive clinical data and medical history of autistic subjects and their first-degree relatives have also been stored in the database. The goal of the project is to build a biorepository that interacts with a database of extensive clinical information collected electronically at each recruiting sites, enabling genetic, genomic and proteomic research on ASD. Although the target research areas are multidisciplinary (from genetic and biomarker research, to epidemiological studies), the design of the project was driven mainly by its genetic goals. The project aims at collecting small families formed by a proband (ASD child), a sibling when available, and their parents. The collection of families would allow performing family-based genetic association studies as well as the screening of genetic variants in cases and affected and unaffected first-degree relatives.

### Recruiting sites and Ethical issues

The biorepository is hosted by the Department of Neurosciences, Biomedicine and Movement Sciences at the University of Verona, Italy. Recruitment has begun in 2008 and has been conducted thus far at thirteen different Italian sites (Verona, Acireale (Catania), Bari, Bologna, Brescia, Cagliari, Lecco, Napoli, Padova, Pisa, Rimini, Roma, and Troina (Enna).

Approximately half (48%) of the patients recruited were referred by the primary care physician, 31% by a tertiary care center, and the rest by schools, or outpatient clinics.

A study protocol for the clinical assessment and sample management was written and agreed by the main centers and founding members of the network. Candidate recruiting sites were selected among clinical centers with a demonstrated ability to assess autism families as described in the project protocol. The study and its protocol were approved by the Azienda Ospedaliera Universitaria Integrata of Verona Ethic Review Board. Furthermore, each recruiting site obtained independent approval to conduct the project by their respective local ethical review committees. Eighteen sites joined the Italian Autism Network, with thirteen actively recruiting. All adult subjects participating in this project gave their consent (or the consent for their chidren) to donate biological samples and clinical and demographic information to participate in this study; assent to participate to this study from the children was obtained whenever possible.

### Subjects

Cases, parents and, whenever available, sibling and/or non-affected individuals identified as controls, were recruited. Based on the selection criteria and the family composition, different type of families can be summarized through the pedigree analysis. We used the R package *kinship2* with the pedigree shrinking function to gain a metric of pedigree structure in terms of *bitSize* (a measure is defined as 2 * # NonFounders - # Founders, see [31]).

### Inclusion and Exclusion Criteria

A proband child or affected sibling was eligible for recruitment when all the following inclusion criteria apply: 1) at least four years of age at the time of the interview, 2) having at least one parent or legal guardian giving voluntary written consent for their children to take part to the research project, 3) meeting DSM-IV criteria for Autistic Disorder or Asperger’s Disorder, or Pervasive Developmental Disorder Not Otherwise Specified (PDD-NOS), 4) reaching score cut-off in ADI-R. Siblings of the proband were recruited when available, and defined as unaffected, if they met the following criteria: 1) at least 4 years of age at the time of recruitment, 2) devoid of any comorbid psychiatric illness as determined by: i) the Child Behavioural Checklist (CBCL) or by the Kiddie Schedule for Affective Disorders and Schizophrenia for children (Kiddie-SADS), ii) absence of social skills impairment as determined by the Social Responsiveness Scale (SRS), iii) the Broader Phenotype Autism Symptom Scale (BPASS) (administered to evaluate the presence of autism spectrum symptoms). Individuals were not eligible for recruitment if any of the following criteria apply: 1) history of serious head injury, encephalitis or tumors, 2) age above 18; 3) Diagnoses of Childhood Disintegrative Disorder or Rett Syndrome, diagnosis of known ASD-related genetic syndromes or presence of severe mental retardation (IQ < 20) that would compromise the validity of the diagnosis of autism.

### Clinical assessment

Diagnosis was established by experienced child psychiatrists after completing full neurological, psychiatric and physical examination, and after reviewing and completing the Autism Diagnostic Interview - Revised (ADI-R) and the Autism Diagnostic Observation Schedule (ADOS) that were administered by trained and certified clinicians. The cognitive level of children was assessed whenever possible through either one of the following scales: Wechsler Intelligence Scale for Children, Wechsler Preschool and Primary Scale of Intelligence, Leiter International Performance Scale or Griffiths Mental Developmental Scale. Additional structured evaluation included the Children’s Global Assessment Scale (CGAS) and, for subjects with suspect of Asperger’s Disorder, the Krug Asperger’s Disorder Index (KADI). The KADI is an individually administered, norm-referenced screening instrument that provides useful information for determining the diagnosis of Asperger’s Disorder. It is standardized for use with individuals 6 through 21 years of age. Instrumental and laboratory examination of the fragile X (FRAX) expansions and major cytogenetic abnormalities were conducted for all the autism individuals and their siblings. MRI scan and awake and nap EEG were evaluated in all patients, if they were not available from previous assessments conducted at a close time.

Parents of the probands and siblings of age 18 and above were assessed as follows: i) Cognitive level: four items of the WAIS (arithmetic, vocabulary, block design, picture arrangement); ii) Language: Token test; iii) Autism spectrum symptoms: BPASS Broader Phenotype Autism Symptom Scale (BPASS); iv) Psychopathology: SCID screening interview.

Non-affected siblings of less and 18 years of age recruited were assessed with medical, psychiatric and neurologic examination. For each non-affected sibling, cognitive levels are established via the four items of the Wechsler Intelligence Scale for Children (WISC) that has shown to produce a reliable measure of I.Q. [32]. The four items include the arithmetic, vocabulary, block design, and picture arrangement. Language competence was assessed with the Peabody Picture Vocabulary test (PPVT-R), for ages between 4 and 12, and the Token Test for age between 12 and 18. Social Responsiveness Scale (SRS) was also administered to evaluate socialization. Psychopathology in non-affected siblings was assessed via the Child Behavioural Checklist (CBCL). The following definitions were used to define the presence of psychopathology: T-score: normal <65; at risk: 66–74; pathological: >75 (2.5 SD from normal average) if scores less above 75 (presence of psychopathology) the Kiddie-SADS interview is administered to diagnose the psychiatric interview.

### Sample collection, handling and storage

The Italian Autism Network collected DNA, plasma, RNA and EBV transformed lymphocytes in the biorepository from each of the individuals assessed. From all individuals eligible to participate in this project (as either cases or unaffected relatives) for whom a signed informed consent was obtained, the following peripheral blood samples were taken: 1) 2× 6-mL in ACD blood tubes for DNA extraction; 2) 1× 6mL ACD tube for extracting lymphocytes to be transformed with Epstein Barr virus (EBV) in LCLs; 3) 1× 6-mL in EDTA tube to obtain plasma aliquots for proteomic analysis, and 4) 2× 2.5-mL in PAX gene tubes for obtaining RNA. Plasma aliquots (0.2 mL) were prepared by centrifugation of EDTA tubes and stored at −80°. Blood-derived samples for RNA extraction were prepared by gently mixing

PAX gene tubes, storing them for 2 hours at room temperature, cooling them at −20° and later at −80° to be ready for shipment. Samples for DNA extraction and cell lines transformation were shipped at room temperature to the biorepository on the same day of collection. Frozen samples (plasma, blood samples for RNA) were shipped via express courier in dry ice to the biorepository.

Upon delivery at the biorepository, Plasma aliquots and PAX gene tubes were stored at -80°. DNA extraction was performed using the Puregene Blood Kit (Gentra Systems, Minneapolis, MN, US), a modified salting-out precipitation method, following the manufacturer’s instructions. Each DNA sample was then quality controlled and quantified using NanoDrop ND 1000 spectrophotometer (Thermo Scientific, Wilmington, DE, US). RNA samples from lymphocytes were isolated with QIAcube system, quantified with NanoDrop and analyzed for quality control by use of the RNA 6000 NanoLabChip (Agilent Technologies, Santa Clara, CA, US). The biorepository also transformed the lymphocytes with EBV into lines that were and then stored in Liquid Nitrogen. All relevant data regarding the samples are entered into a computerized system that allows matching the information of the samples with the clinical data stored in the centralized clinical database.

### Data security procedure for the clinical database

All blood, plasma, cell lines, DNA and RNA samples are coded and stored under secure conditions with restricted access according to the Italian privacy policies required for keeping secure sensitive data. The clinical information collected at each site is entered into an encrypted system and stored in a database. Codes corresponding to the research subjects are known only by the clinician responsible for the research project, who obtained the informed consent from the parents of affected subjects and any sibling at each recruiting site (Site Principal Investigator), and by the clinical personnel directly involved in the assessment of the research subject, as required by standard clinical practice.

## Discussion

The Italian Autism Network (ITAN) was initiated under the sponsorship of the Fondazione Smith Kline with the aim of creating an open resource for clinical, genetics and biomarker research in autism. Here we described the protocol used to build up this novel family collection and the associated database and biorepositorynk. Each family is composed by at least one ASD affected child and, when possible, a discordant sibling.

### Recruitment progress to date

To date, the ITAN consortium has recruited 812 subjects, i.e. 252 probands and 560 first degree relatives, for a total of 249 families. In Table 1 we provide information on the distribution of samples among recruiting centers and the demographic of the collection. In Figure 1 we illustrate the distribution of pedigree structures among families recruited in the study.

**Figure 1.**
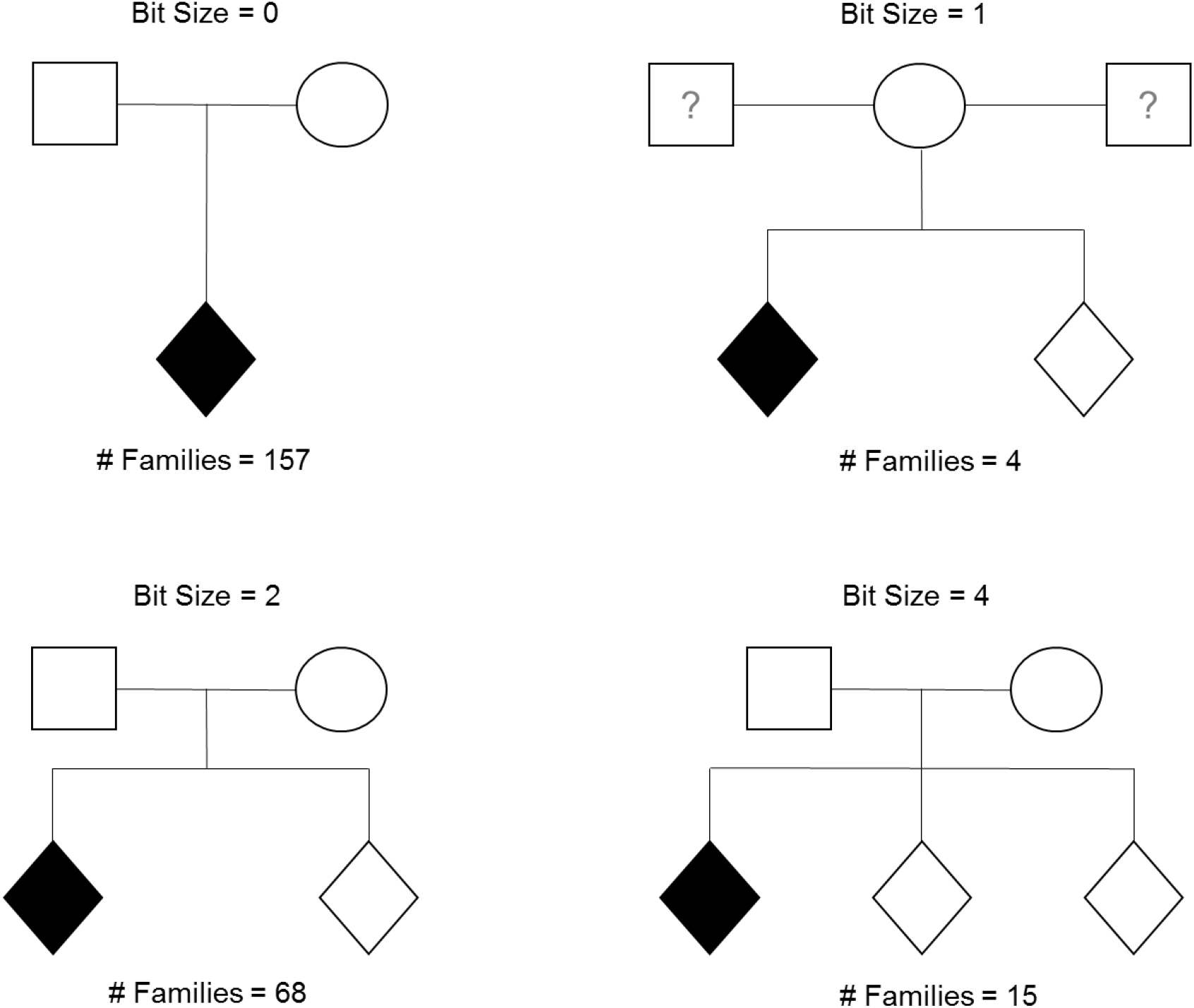
Pedigree structures of families recruited into the ITAN collection based on the bitSize measure (only most represented pedigree structures are shown). For each pedigree structure, the number of families of the ITAN dataset are reported.

**Table 1.** Distribution of subjects among recruiting centers and summary of demographic information.

| Site | Families | Parents | Unaffected siblings | Autism Spectrum Disorders |  |  |  |  |  |  |  |  |
| --- | --- | --- | --- | --- | --- | --- | --- | --- | --- | --- | --- | --- |
|  |  |  |  | ASD cases | Gender (% males) | Average Age (SD) | Ethnicity |  |  | Diagnosis |  |  |
|  |  |  |  |  |  |  | AFR | OTH | CEU | AD | ASPG | PDD-NOS |
| 3 | 66<sup>§</sup> | 135 | 57 | 68 | 85.29 | 9.44 (3.84) | 8 | 0 | 60 | 50 | 9 | 9 |
| 4 | 4<sup>§</sup> | 8 | 0 | 5 | 100.00 | 10.60 (1.67) | 0 | 0 | 5 | 3 | 0 | 2 |
| 5 | 27* | 54 | 1 | 26 | 92.31 | 5.38 (2.17) | 0 | 2 | 24 | 18 | 0 | 8 |
| 6 | 21<sup>§</sup> | 42 | 7 | 22 | 72.73 | 9.45 (4.68) | 11 | 0 | 11 | 18 | 1 | 3 |
| 7 | 17 | 34 | 10 | 17 | 82.35 | 10.71 (4.55) | 8 | 3 | 6 | 13 | 3 | 1 |
| 8 | 32* | 63 | 0 | 31 | 83.87 | 6.71 (2.77) | 16 | 0 | 15 | 5 | 2 | 24 |
| 9 | 17 | 33 | 11 | 17 | 88.24 | 11.24 (4.12) | 0 | 0 | 17 | 11 | 1 | 5 |
| 10 | 8<sup>§</sup> | 16 | 1 | 9 | 88.89 | 7.89 (3.95) | 2 | 0 | 7 | 6 | 0 | 3 |
| 11 | 13 | 27 | 2 | 13 | 76.92 | 11.77 (4.55) | 4 | 0 | 9 | 4 | 6 | 3 |
| 12 | 11<sup>§</sup> | 22 | 3 | 12 | 75.00 | 5.25 (1.71) | 0 | 0 | 12 | 4 | 0 | 8 |
| 13 | 9* | 17 | 2 | 8 | 87.50 | 10.88 (2.85) | 0 | 1 | 7 | 5 | 1 | 2 |
| 16 | 4 | 8 | 3 | 4 | 100.00 | 10.50 (5.51) | 0 | 0 | 4 | 4 | 0 | 0 |
| 17 | 20 | 40 | 2 | 20 | 95.00 | 10.05 (4.67) | 2 | 0 | 18 | 12 | 2 | 6 |
| Tot | 249 | 499 | 99 | 252 | 86.78 | 9.22 (3.62) | 51 | 6 | 195 | 153 | 25 | 74 |

Legend: Distribution of subjects among recruiting centers and summary of demographic information. \*indicates centers where family recruitment is not completed (*eg* affected subjects less than N of families). <sup>§</sup>Indicates centers which have recruited families with >1 affected subjects

### Clinical phenotypes

The clinical assessment was derived by full neuropsychiatric evaluation and structured interviews (i.e.: ADI-R, and ADOS) by appropriately trained and certified staff. The average age at interview of the affected and non-affected subjects recruited to date is 8.89 and 11.53, respectively, while the ratio of males in the affected samples is 85.31%, reflecting the higher incidence of autism in males reported for the disease in the general population. For the non-affected siblings, the proportion of males is 55.55%. Figure 2 reports the distribution of the cognitive level for each diagnostic category (DSM IV), as measured by I.Q. assessment. As expected, high functioning Asperger subjects have higher values of I.Q. compared to PDD-NOS and AD subject group (the latter showing the lowest I.Q. values). Only 7.1% of the affected subjects (n= 18) have serious mental retardation with a low I.Q. (<35) as shown in Tab. 2. It should be however noted, that I.Q. below 20 was an exclusion criterion for the probands.

**Figure 2:**
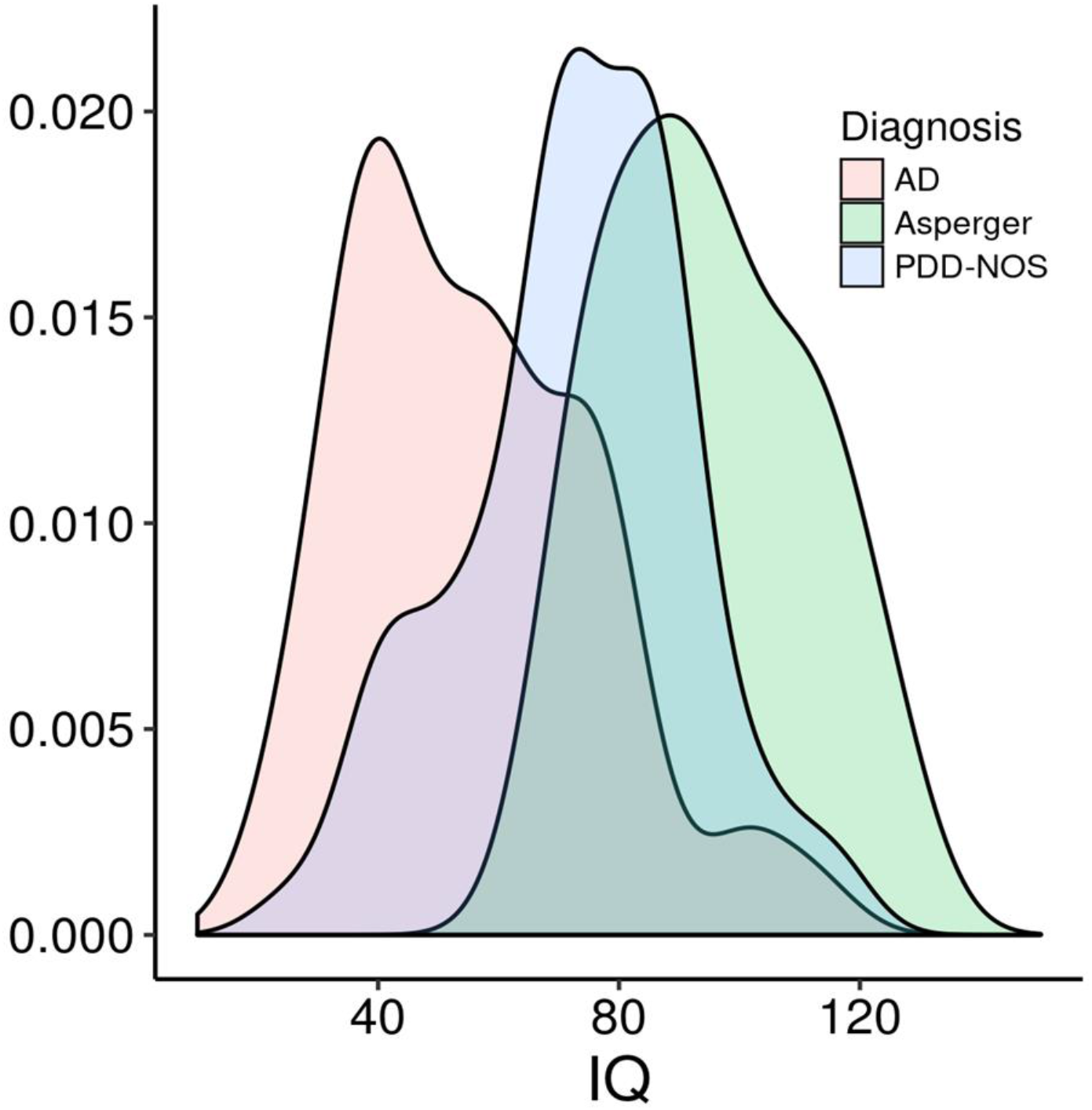
distribution of the cognitive level of ASD subjects of the ITAN collection, as measured by IQ assessments, for each of the three diagnostic categories.

**Table 2.** Intellectual quotients (I.Q.) of affected subjects across diagnostic categories

| **Diagnosis** | **IQ<35** | **IQ= 35-50** | **IQ=50-75** | **IQ>75** | **not available** |
| --- | --- | --- | --- | --- | --- |
| Autistic Disorders | 17 | 53 | 49 | 29 | 5 |
| Asperger Syndrome | 0 | 0 | 4 | 20 | 1 |
| Pervasive Developmental Disorders | 1 | 9 | 29 | 33 | 2 |

### Preliminary genetic investigations

So far, DNA samples from subjects of the ITAN collection have been utilized for testing candidate risk variants for autism derived from large GWAS investigations [26,27,33]. Furthermore, the regulation of candidate genes has been studied by analyzing RNA expression in LCLs from selected probands and non-affected siblings [34]. A large proportion of the discordant sibling pairs (for a total of 152 samples) was genotyped on the PsychArray (Illumina Inc., San Diego, California, USA) and blood RNA was sequenced on the Illumina HiSeq 2000 RNA Seq Platform. The availability of both genotype and RNA expression data on a genome scale is being exploited for a novel integrative approach aimed at the identification of genetical-genomic biomarker signatures for autism [35,36].

In conclusion, the ITAN collection clearly represents a valuable resource for the identification and characterization of common and rare genetic risk variants for autism, thanks to the accuracy of the clinical assessment and the family design. The availability of different types of biological samples makes it a unique collection for the conduct of biomarkers investigations integrating multiple “omic” platforms.

**Abbreviations**
AD: Autistic Disorder
ADI-R: Autism Diagnostic Interview-Revised
ADOS: Autism Diagnostic Observation Schedule
ASD: Autism Spectrum Disorder
BPASS: Broader Phenotype Autism Symptom Scale
CBCL: Child Behavioural Checklist
CGAS: Children’s Global Assessment Scale
DSM-IV: Diagnostic and Statistical Manual of Mental Disorders, 4TH Edition, Text Revision
EEG: Electroencephalography
FRAX: Fragile X
GWAS: Genome-Wide Association Study
IQ: Intelligent Quotient
ITAN: Italian Autism Network
KADI: Krug Asperger’s Disorder Index
Kiddie-SADS: Kiddie Schedule for Affective Disorders and Schizophrenia for children
LCL: Lymphoblastoid Cell Lines
PDD-NOS: Pervasive Developmental Disorder Not Otherwise Specified
PPVT-R: Peabody Picture Vocabulary Test - Revised
SRS: Social Responsiveness Scale
WAIS: Wechsler Adult Intelligence Scale
WISC: Wechsler Intelligence Scale for Children

### Declarations

#### Ethics approval and consent to participate

The study protocol was in first instance approved by the Verona Hospital Ethical Review board (study protocol AUT-SFK001, CE1419), and subsequently by the Ethical Review Committees of each recruiting site. A multidisciplinary Scientific and Ethical Committee of the Fondazione SmithKline, “Area Ricerca Autismo”, i) created the protocol; ii) ensured that the use of the samples and clinical database was consistent with what consented by the research participants; iii) ensured that research was conducted with sound methodology and by accredited institutions. The Scientific and Ethic committee was chaired by Prof. Bernardo Dalla Bernardina and composed by (in alphabetical order) the Secretary Lucio Da Ros (formerly at GSK), Luciano Eusebi (Università Cattolica di Milano), Gabriele Masi (Università di Pisa), Pierandrea Muglia (UCB Pharma Belgium), Franco Nardocci (Ospedale Infermi di Rimini), Donata Pagetti Vivanti (President Autismo Italia e Autismo Europa), Pier Franco Pignatti (Università di Verona), Mons. Sergio Pintor (Vescovo di Ozieri (SS), Director of Uff. Naz. CEI Pastorale per la Sanità), Emiliangelo Ratti (formerly at GSK), Paolo Rizzini (Vice Presidente Fondazione Smith Kline), Alessandro Zuddas (Università di Cagliari).

#### Availability of data and material

The ITAN is an open resource. In 2012 the Fondazione Smith Kline transferred the ownership to a newly created not-for-profit organization ‘Italian Autism Network’ Foundation that govern the use of the resource with appropriate governance (see [37] for details). To gain data access, interested researchers should submit an official request to the Foundation. Data will be released only for specific research purposes, upon a positive assessment of a project proposal by the ITAN Scientific Committee.

#### Authors’ contributions

PM, LDR, PR, BDB, ER designed the study, wrote the protocol and coordinated the implementation of the network, its infrastructure and logistics.

ET has responsibility for the biorepository and was actively involved in writing the sample collection and shipping guidelines.

MF, TKT, LDR, PM and ED wrote the manuscript.

BDB, ET, FN, AZ, ER, PM, ED are members of the Scientific Committee, reviewed and approved the manuscript.

## Acknowledgments

We would like to thank first of all family members who provided their time and samples that made this collection possible.

We would also like to thank the following individual who contributed to the creation of ITAN:

Leonardo Zoccante, Flavio Boscaini. Servizio di Neuropsichiatria Infantile Azienda Ospedaliera Istituti Ospitalieri di Verona Policlinico G.B. Rossi, Verona.

Maurizio Elia, Giuseppa Di Vita, IRCCS “Associazione Oasi Maria SS.” (Istituto di Diritto Privato), Troina (Enna).

Massimo Molteni, Elisa Mani, Elisa Ceppi, IRCCS “Eugenio Medea”, Monza.

Antonio Pascotto, Carmela Bravaccio Neuropsichiatria Infantile - Seconda Università degli Studi di Napoli.

Alessandra Tiberti, Filippo Gitti, Giovanni Allibrio, Azienda Ospedaliera Spedali Civili di Brescia. Brescia.

Lucia Margari, Anna Linda Lamanna, Andrea DeGiacomo, Unità Operativa di Neuropsichiatria Infantile Dipartimento di Scienze Neurologiche e Psichiatriche Università degli Studi Bari, Bari.

Alessandro Zuddas, Laura Anchisi, Università degli Studi di Cagliari - Centro per lo Studio delle Terapie Farmacologiche in Neuropsichiatria dell’infanzia e dell’adolescenza, Cagliari.

Paolo Curatolo, Barbara Manzi, Arianna Benvenuto, Clinica S.Alessandro - Cattedra di Neuropsichiatria Infantile - Università “Tor Vergata” - Policlinico Tor Vergata, Roma. Gabriele Masi, Raffaella Tancredi, IRCCS Stella Maris, Pisa.

Franco Nardocci, Serenella Grittani, Tiziana Piroddi, Milena Andruccioli, Ospedale Infermi - Divisione Neuropsichiatria Infantile - Centro per l’autismo, Rimini.

For the biorepository we thank Pier Franco Pignatti, Elisabetta Trabetti, Paola Prandini.

For the executive group we thank Diego Cosentino, Sonia Colcera (GSK), Maurizio Bassi (Fondazione Smith Kline), Luigi Napolitano (Fondazione Smith Kline), Francesca Arzone (GSK), and Ugo Bianchi and Maurizio Martignano from Ad-hoc Sistemi. We would like to acknowledge the multidisciplinary Scientific Committee of the

Fondazione SmithKline “Area Ricerca Autismo” that guided the project since its conception and regularly monitored its progression for ethical and scientific rigor.

^§^ITAN (the Italian Autism Network) is composed by: Giovanni, Alibrio; Laura, Anchisi; Milena, Andruccioli; Arianna, Benvenuto; Pier Antonio, Battistella; Flavio, Boscaini; Carmela, Bravaccio; Elisa, Ceppi; Diego, Cosentino; Paolo, Curatolo; Lucio, Da Ros; Bernardo, Dalla Bernardina; Andrea, De Giacomo; Giuseppa, Di Vita; Enrico, Domenici; Massimo, Elia; Filippo, Gitti; Serenella, Grittani; Anna Linda, Lamanna; Elisa, Mani; Barbara, Manzi; Lucia, Margari; Gabriele, Masi; Massimo, Molteni; Pierandrea, Muglia; Franco, Nardocci; Antonio, Pascotto; Antonia, Parmeggiani; Pier Franco, Pignatti; Tiziana, Piroddi; Paola, Prandini; Emiliangelo, Ratti; Paolo, Rizzini; Sebastiano, Russo; Renato, Scifo; Raffaella, Tancredi; Alessandra, Tiberti; Elisabetta, Trabetti; Leonardo, Zoccante; Alessandro, Zuddas.

